# Beyond Differential Expression: Embracing Cell-to-Cell Variability in Single-Cell Gene Expression Data Analysis

**DOI:** 10.1101/2024.08.08.607086

**Authors:** Victoria Gatlin, Shreyan Gupta, Selim Romero, Robert S. Chapkin, James J. Cai

## Abstract

Single-cell RNA sequencing (scRNA-seq) has revolutionized our understanding of cellular variability by capturing gene expression profiles of individual cells. The importance of cell-to-cell variability in determining and shaping cell function has been widely appreciated. Nevertheless, differential expression (DE) analysis remains a cornerstone method in analytic practice, by focusing exclusively on mean expressional differences, current computational analyses overlook the rich information encoded by variability within the single-cell gene expression data. Thus, there is a need for more relevant approaches to assess data variability rather than the mean in order to offer a deeper understanding of cellular systems. Here we present spline-DV, a novel statistical framework for differential variability (DV) analysis using scRNA-seq data. The spline-DV method identifies genes exhibiting significantly increased or decreased expression variability among cells derived from two experimental conditions. Case studies show that DV genes identified using spline-DV are representative, and functionally relevant in the tested cellular conditions including obesity, fibrosis, and cancer.

## Introduction

It is well established that transcription of many genes occurs in bursts (Raj, Peskin, Tranchina, Vargas, & Tyagi, 2006; Suter et al., 2011), resulting in large mRNA number variability between cells, even if they are from the same underlying state. The importance of cell-to-cell variability of gene expression in determining and shaping cell function has been recognized for some time (Altschuler & Wu, 2010; Dueck, Eberwine, & Kim, 2016; Losick & Desplan, 2008). Increased variability in gene expression has been associated with the differentiation of embryonic stem cells (Stumpf et al., 2017), reprogramming of induced pluripotent stem cells (Buganim et al., 2012), circadian rhythm regulation (Droin et al., 2021; Phillips et al., 2021), differentiation (Mojtahedi et al., 2016; Richard et al., 2016), and aging (Bahar et al., 2006; Martinez-Jimenez et al., 2017). In this single-cell omics era, investigators now have a higher resolution tool such as single-cell RNA sequencing (scRNA-seq) to reveal gene expression variability between cells, allowing us to better understand its role. In a recent study (Osorio et al., 2019), we demonstrate that single-cell gene expression variability is intrinsically linked to the function of genes.

Individual cells of the same cell type exhibit stochastic gene expression that goes beyond technical noise, and this variability is primarily driven by highly variable genes (HVGs), which may not be highly expressed in all cells, but show significant variation between cells and predominantly contribute to cell-type-specific functions. If the variability is indeed central to cellular function, as our findings suggest, the development of analytical methods and computational tools for single-cell data should take this variability-centric view into account. Unfortunately, in practice, the importance of variability is often overlooked. Thus, current methods developed under non-variability-centric principles may require significant refinement. For example, many single-cell data analytical frameworks are mean-centric and neglect gene expression variability by design.

Differential expression (DE) analysis has been a mainstay of gene expression studies since the days of microarray and bulk RNA-seq. Traditionally, DE analysis focuses on identifying genes that are up- or down-regulated (increased or decreased expression) between conditions, typically employing a basic mean-difference approach. This approach is also commonly applied to scRNA-seq data, with popular R packages like DESeq2, EdgeR, and limma being directly adapted for this purpose (Luecken & Theis, 2019). While many DE methods have been specifically developed for scRNA-seq data (Das, Rai, & Rai, 2022), these single-cell specific DE methods do not outperform their bulk counterparts (Van den Berge et al., 2018). These methods often consider cell-to-cell variability and dropout as technical noise, overlooking the fact that both the variability and dropout contain important biological information (Bouland, Mahfouz, & Reinders, 2023; Qiu, 2020). Additionally, existing DE methods show inconsistency in identifying DE genes, again suggesting that they fail to capture the full spectrum of biologically relevant expression changes (T. Wang, Li, Nelson, & Nabavi, 2019).

Here we argue against methods focused solely on detecting mean expression differences in scRNA-seq data. We propose a paradigm shift that acknowledges the central role of gene expression variability in cellular function and challenges the current dominance of mean-based DE analysis in single-cell studies. This shift becomes particularly urgent in light of recent efforts, such as the highly cited study by (Squair et al., 2021), which proposes analyzing pooled, pseudobulk scRNA-seq data to mimic bulk RNA-seq for DE analysis. While such approaches may seem appealing, they disregard the primary strength of scRNA-seq—its ability to capture cell-to-cell variability. Critically, (Squair et al., 2021) use bulk RNA-seq DE results as a reference to evaluate the performance of scRNA-seq DE methods, reinforcing a “mean-centric” view that considers bulk RNA-seq as the gold standard despite its limitations in capturing cellular variability. Thus, the assumption that the pseudobulk method perfectly recovers bulk RNA-seq expression patterns needs further investigation, as these techniques likely reveal distinct biological phenomena. It is important to be aware of the prevalence of “mean-centric thinking” in molecular biology, where microarray and bulk RNA-seq analysis methods dominate scRNA-seq data analysis. We suggest that scRNA-seq data analysis should embrace the role of inherent gene expression variability in defining cellular function and move beyond mean-based approaches.

## Results

### Illustrating the “variation-is-function” concept

(Dueck et al., 2016) proposed the “variation-is-function” hypothesis suggesting that cell-to-cell gene expression variability plays a significant role in manifesting population-level cellular function. Our previous study of scRNA-seq data supports this hypothesis (Osorio et al., 2019). We detected highly variable genes (HVGs) in homogenous cellular populations and found that, for each different cell type, the functions of HVGs are enriched with biological processes and molecular functions precisely relevant to the biology of the corresponding cell type. **Fig. 1** illustrates this “variation-is-function” concept. Three summary statistics—mean, coefficient of variation (CV), dropout rate—was used to depict each gene’s expression across cells. A spline fit algorithm introduced in the scGEAToolbox (Cai, 2019) was used to build a spline-fit curve going through all genes. HVGs were identified as those that deviate most from the spline-fit curve. In the simulated data (**Fig. 1A**), there were no HVGs as all genes align closely along the spline-fit curve. Undifferentiated cells tended to have fewer detected HVGs and less gene expression variability compared to differentiated cells, as shown by the human embryonic stem cells (hESCs, **Fig. 1B**) vs. human umbilical vein endothelial cells (HUVECs, **Fig. 1C**). Overall, these results suggest that gene expression variability increased as cells differentiate. As undifferentiated cells (e.g., hESCs) differentiate into more distinct, mature forms (e.g., HUVECs), additional genes are expressed variably and become HVGs. One such gene is *ANKRD1* (shown in **Fig. 1C**), which encodes ankyrin repeat domain 1 transcription factor that is localized to the nucleus of endothelial cells.

**Fig. 1.**
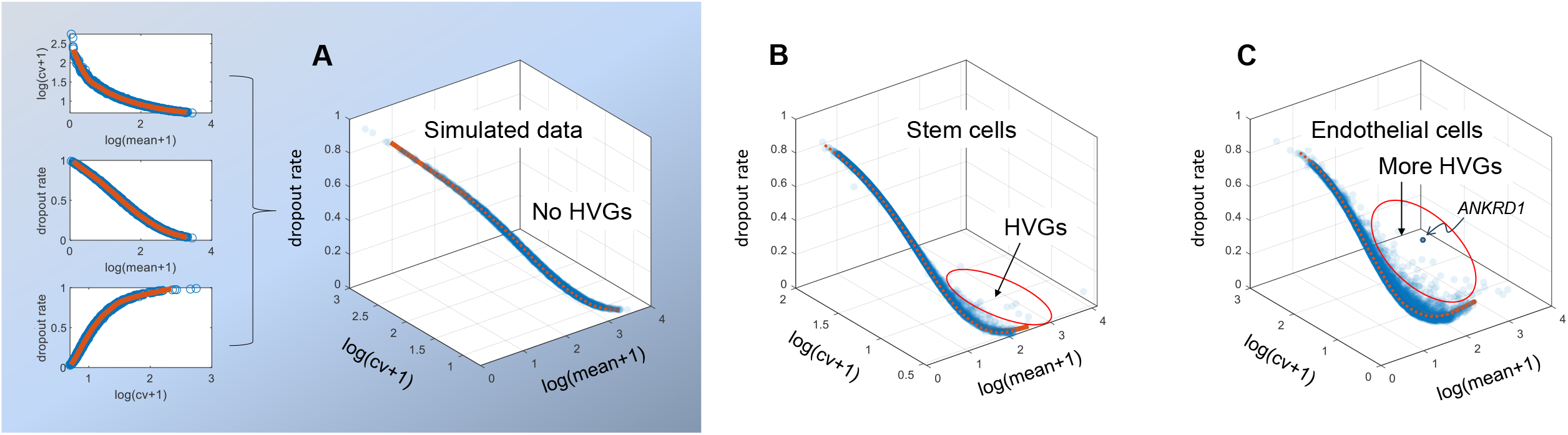
Single-cell gene expression variability, driven by highly variable genes (HVGs), increases with cell differentiation level. (**A**) Simulated data. The data is a 5,000-by-3,000 (gene-by-cell) count matrix, which was randomly generated using a negative binomial (NB) model with a fixed dispersion level of 0.1 (Lun, Bach, & Marioni, 2016). Three 2D-scatter plots show the pairwise relationships between three gene-level metrics: mean, CV, and dropout rate. These metrics are combined into a 3D-scatter plot, where genes (blue dots) are distributed along a characteristic spline-fit curve (red line). The position of each gene is determined by its mean, CV, and dropout rate. (**B**) Human embryonic stem cell (hESC) data generated using scRNA-seq in the study of (Xu et al., 2023). Similar to **A**, gene expression variability is shown with a 3D-scatter plot of genes (blue dots). A spline-fit curve (red line) captures the global pattern of the three metrics across all genes. (**C**) Human umbilical vein endothelial cell (HUVEC) data representing a differentiated cell type. In contrast to undifferentiated hESCs in **B**, HUVECs show more HVGs and higher overall single-cell gene expression variability. As an example, one of the HVGs, *ANKRD1*, is indicated in the 3D-scatter plot.

### Identification of differentially variable genes

We introduce the spline-DV—a nonparametric, model-free method for analyzing changes in single-cell gene expression variability between two conditions. The goal is to identify differentially variable (DV) genes. These genes are supposed to be functionally more engaged or transcriptionally more active in one condition than the other. Analyzing the variability changes and detecting DV genes have the potential to provide a more comprehensive understanding of the transition of active cell state across different conditions. **Fig. 2** illustrates the spline-DV method. This method accounts for mean expression, CV, and dropout rate statistics in a 3D model, with these being the x, y, and z coordinates, respectively. Within this 3D space, it creates a spline-fit curve for each condition (**Fig. 2A**). The two spline curves, visualized in blue and red, are then projected into the same 3D space (**Fig. 2B**), allowing for a comparative assessment across conditions. For a given gene (e.g., Gene A), the position of the gene in the 3D space was determined by the observed expression profile values of the gene (i.e., mean expression, CV, and dropout rate). From the position to the nearest point on the spline-fit curve, we created a vector and let the vector originate from the nearest point. For the same gene (i.e., Gene A), two vectors were created for two conditions, 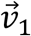 and 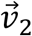, respectively. The two vectors represent the deviation of the same gene (i.e., Gene A) away from its expected positions with respect to the two conditions. The magnitude of each vector was used to quantify the level of deviation, i.e., the expression variability of the gene in each condition. To compute the level of differential variability between two conditions, the difference between 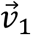 and 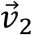 was computed (**Fig. 2C**). The resulting DV vector, 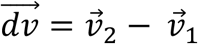, captured the difference in variability between two conditions. In this way, a fair comparison between conditions was made by first comparing each gene to its expected statistics rather than making a direct comparison. The *DV score* defined as the magnitude of 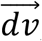, measured the level of differential variability of the gene between two conditions (**Methods**). Finally, spline-DV ranked the list of genes based on their DV scores. Top DV genes across conditions were prioritized for further investigation as they are likely to play crucial roles in the biological processes under study.

**Fig. 2.**
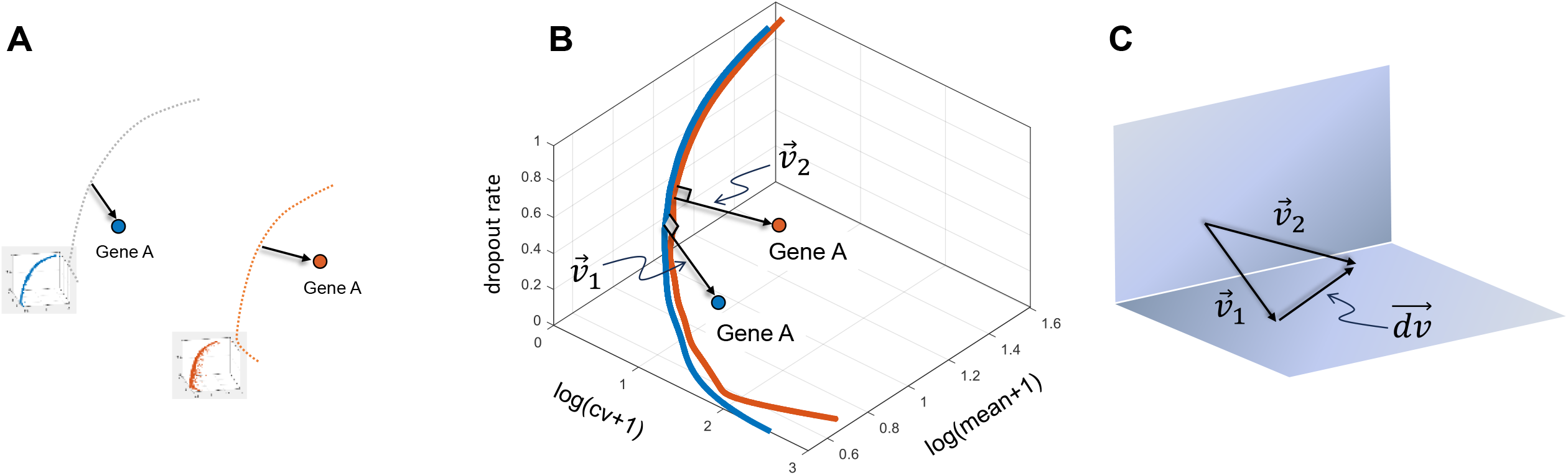
Illustration of the spline-DV method proposed in this study. (**A**) Two spline-fit curves are created independently with scRNA-seq data of two conditions, blue for condition 1 and red for condition 2. These spline curves represent the “expected” expression profile for genes based on their statistical properties in each condition. (**B**) Two spline-fit curves are projected into the same 3D space for comparison. For a given gene, Gene A, its position is compared with respect to the condition 1 spline-fit curve. The vector 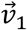 extends from the closest point on the blue curve (spline-fit) to the blue point (Gene A in condition 1). Similarly, for condition 2 (red point), the vector 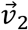 extends from the closest point on the red curve to the red point (Gene A in condition 2). The L2-norm of 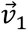 (or 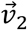) quantifies the deviation of the same Gene A from its expectation in condition 1 (or condition 2). (**C**) Given that the deviation is the shortest Euclidean distance from the blue (or red) point to the blue (or red) spline-fit curve, representing the level of expression variability of the gene in condition 1 (or condition 2), 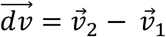, and the L2-norm of 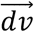, i.e., DV score, is used to measure the level of differential variability.

### Case studies

#### Case study 1 – More insightful genes identified in adipocytes in response to diet-induced obesity

We applied the spline-DV method to real scRNA-seq data sets to assess its performance. Our first case study utilized a scRNA-seq data set generated from a study focusing on diet-induced obesity (Sarvari et al., 2021)(**Fig. 3A**). Adipocytes dissociated from adipose tissues in mice fed either low-fat diet (LFD) or high-fat diet (HFD) for 18 weeks were collected and compared (**Fig. 3B**). Using the spline-DV method, we identified DV genes exhibiting differential variability between two conditions, e.g., *Blcap, Nnat*, and *Lyz2* (**Fig. 3C**). In addition, *Plpp1*, which encodes a protein of the phosphatidic acid phosphatase (PAP) family, with increased variability in the HFD sample, and *Thrsp*, which encodes thyroid hormone-inducible hepatic protein exhibited a decreased variability in the HFD sample (**Fig. 3D**). Previous studies show that *Plpp1* deletion increases endogenous lipid lysophosphatidic acid concentrations in hepatocytes and reduces hepatocyte glucose production (Taddeo et al., 2017), and *Thrsp* deletion decreases mitochondrial respiration and fatty acid oxidation, contributing to metabolic dysfunction in obese adipose tissue (Ahonen et al., 2022). Interestingly, DV genes collectively are enriched with the core adipocyte pathways related to *lipid metabolism, insulin response, and biosynthesis*. (Sarvari et al., 2021) identified the *fatty acid biosynthesis* pathway using DE analysis. We detected the same pathway using spline-DV. Beyond this, using spline-DV, we exclusively identified a DV gene—*HADH*, which was not identified as a DE gene. *HADH* encodes hydroxyacyl-CoA dehydrogenase, which regulates *fatty acid breakdown* and *tryptophan metabolism* (**Fig. 3E**). Overlap between DE and DV genes is limited (**Fig. 3F**), reflecting the discrepancy between DE and DV methods. Collectively, these findings demonstrate that spline-DV method may afford additional insight into gene functions and cellular states between cell samples from two conditions. These changes are brought about by modifications in gene expression variability rather than mean.

**Fig. 3.**
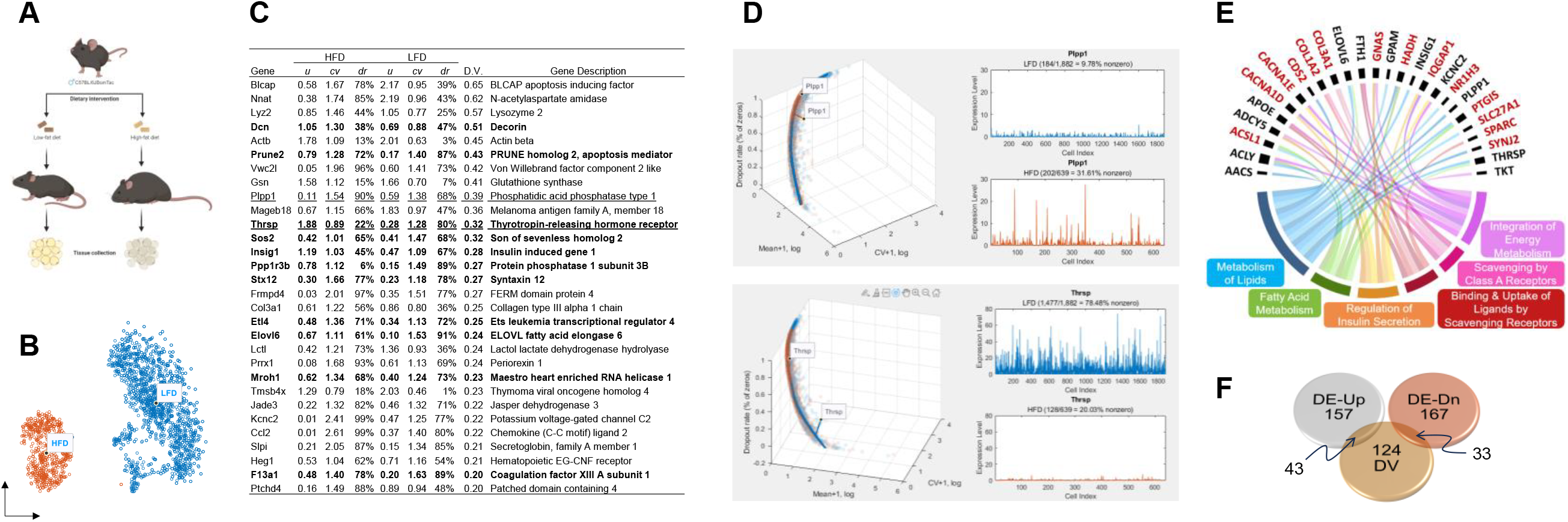
Application of spline-DV method in analyzing scRNA-seq data from a mouse nutrition study. (**A**) Experimental design for the scRNA-seq study (Sarvari et al., 2021) involving low-fat diet (LFD) and high-fat diet (HFD) mice. (**B**) UMAP representation of HFD and LFD adipocyte cells. (**C**) List of top 30 DV genes. Normal font indicates genes with increased variability in HFD samples, while bold font indicates genes with decreased variability in HFD samples. Two genes highlighted with underlined font are selected as representatives. The column D.V. gives the DV score values. (**D**) HFD-LFD spline-DV variance representation of gene statistics (mean, CV, and dropout rate) for the two selected genes, *Plpp1* and *Thrsp*. (**E**) HFD-LFD DV genes of adipocyte cells identifies significant genes and pathways related to the experimental conditions. Black font indicates genes captured by DE and DV, while red font indicates DV genes exclusively. (**F**) Venn diagram of overlapping genes. The top 200 DV genes show small intersections with the top 200 up-regulated and top 200 down-regulated DE genes.

#### Case study 2 – Enhanced knowledge of gene activity in hepatic stellate cells with simulated chronic liver fibrosis

Our second case study for the spline-DV method utilized a scRNA-seq data set generated from a study focusing on potential drivers of liver fibrosis (Dobie et al., 2019). The study was designed with a well-established chronic injury model using carbon tetrachloride administration in mice. Carbon tetrachloride effectively mimics human centrilobular fibrosis, allowing researchers to investigate disease mechanisms. Using this publicly available data set, we compared the healthy control and 6-week chronic injury hepatic stellate cell samples. DV genes identified using the spline-DV method were more representative of genes associated with the progression of fibrosis, in contrast to DE genes that were not. For example, DV genes, *Col1a1, Col3a1, Col6a3*, and *Col8a1*, that are directly linked to fibrosis progression, were not captured by DE. The products of these collagen genes are the major protein component of extracellular matrix that accumulates excessively in fibrosis (Ma et al., 2024). In addition, DV genes that encode enzymes involved in *extracellular matrix remodeling* such as *Mmp2, Mmp14, Adamts4, Adamts5, Timp1*, and *Timp3*, were also identified. Pathway analysis revealed that DV genes were enriched in core fibrotic processes of *collagen fibril organization, extracellular matrix organization*, and *regulation of fibroblast proliferation*.

#### Case study 3 – Deeper understanding of driver genes in epithelial cells affected by colorectal cancer

Our third case study utilized a scRNA-seq data set from a study focusing on the malignant transformation of colorectal cancer (Becker et al., 2022). We compared epithelial cells from unaffected and cancerous patient samples. DV genes proved more beneficial for data analysis than DE genes in detecting genes associated with disease progression. For example, DV genes associated with colorectal cancer progression, such as *MACF1, SMOC2*, and *FGFR2* (Jang et al., 2020; Li et al., 2019; Mohamed et al., 2021; Shvab et al., 2016), were not captured by DE. In addition, pathway analysis revealed DV genes involved in *notch signaling* such as *JAG1, GMDS, SORBS2, and RBPJ*. This is noteworthy, because activation of this pathway enhances colonic epithelial cell survival, thereby contributing to tumorigenesis in colorectal cancer (Suman, Das, Ankem, & Damodaran, 2014). Moreover, high expression of *JAG1* is associated with poor prognosis after surgery for colorectal cancer (Sugiyama et al., 2016). Pathway analysis further revealed that DV genes were enriched in essential regulatory processes *including ERBB signaling, ERK1 and ERK2 cascade, protein kinase activity*, and *cell motility*.

## Discussion

Our goal was to develop a robust method, both biologically and statistically optimized, to identify transcriptomic changes at a single-cell level that extends beyond average gene expression changes. Our solution, the spline-DV method, is an analytical framework for comparative gene expression variability analysis, based on single-cell gene expression mean, CV, and dropout rate. Spline-DV is expected to identify novel, functionally relevant genes that are typically overlooked by traditional DE methods.

Statisticians have been undoubtedly influenced by biologists in developing statistical methods for solving biological questions. DE methods are designed to estimate the mean expression changes as accurately as possible. The strategy often involves complex statistical modeling and individual gene expression distribution fitting to precisely estimate mean. However, focusing solely on mean differences may be misguided if variability in gene expression holds a significant contribution in cellular function. Based on this premise, comparison of mean expression levels between samples would not represent the complete biological complexities of the data. Unfortunately, traditional methods are centered on DE analysis. The effort of developing DV analysis is relatively limited (Eling, Richard, Richardson, Marioni, & Vallejos, 2018; Ho, Stefani, dos Remedios, & Charleston, 2008; Ran & Daye, 2017; K. Wang et al., 2014). It is worth noting that (Liu, Kreimer, & Li, 2023) has previously proposed a method using scRNA-seq data to detect changes in overall variability between two groups of cells. The analysis is focused on assessing variability of cell populations rather than genes, on which our spline-DV method focuses.

A shift in focus from mean expression to gene expression variability is essential for a more comprehensive understanding of cellular function. As a promising research direction, after identifying DV genes, we hope to unlock a deeper understanding of cellular function by focusing on the cause of DV. Indeed, evidence from coupled scRNA-seq and scATAC-seq data (i.e., simultaneous profiling of transcriptome and chromatin accessibility and gene expression from the same cell/nucleus) shows that the expression of HVGs is strongly regulated (Zhong, Cui, Yang, & Cai, 2024). Thus, changes in single-cell gene expression variability may be further evaluated using information from correlated single-cell open chromatin profiling.

The spline-DV method has several advantages in addition to its ability to identify significant genes that could be more functionally relevant to the specific cell type under study. One of the advantages is its robustness against the batch effect of input data, which is a common issue in scRNA-seq analyses. For example, we know that the baseline expression of genes often tends to shift between different biological treatment conditions, resulting in batch effect. The spline-DV is resilient to such bias and provides a fair means to compare scRNA-seq data from two treatment groups. This is because the 3D spline curve, the building block of spline-DV, is computed in a treatment-specific manner, i.e., two conditions are processed independently. Other advantages of the spline-DV method are related to its impressive computational efficiency and its unsupervised methodology. The spline-fit method itself has been considered one of the best methods for feature selection (Sheng & Li, 2021).

## Conclusion

We propose the spline-DV method for comparing changes in expression variability in single-cell data between different conditions. The approach, termed “differential variability”, can potentially provide additional insight into the role of gene function within the continuum of cell state transitions compared to traditional DE analysis. Based on the work presented here, we demonstrate the effectiveness of spline-DV using real scRNA-seq data case studies. By embracing cell-to-cell variability, we can gain a deeper understanding of cellular processes and dynamics within complex tissues.

## Methods

### Single-cell gene expression data

The scRNA-seq data for making **Fig. 1A** was generated through computational simulation using the algorithm of (Lun et al., 2016). The real scRNA-seq data sets used for generating **Fig. 1B,C** were derived from human embryonic stem cells (hESCs) and human umbilical vein endothelial cells (HUVECs), respectively. The hESC data set was obtained from the study of (Xu et al., 2023), which focused on the role of FLI1 in the hESC-EC system. The HUVEC data set was obtained from the GEO database using accession number GSM7511518. This scRNA-seq study focused on the role of vascular endothelial cell growth factor (VEGF), with the data derived from a control sample not treated with human recombinant VEGF-A165. The scRNA-seq data sets used for case studies were from the studies of (Sarvari et al., 2021), (Dobie et al., 2019), and (Becker et al., 2022), respectively.

### The spline-DV framework

The input of the spline-DV is two lists of three statistics: mean, CV, and dropout rate, depicting expression profile of genes across cells in two conditions. The statistics are defined as follows, mean 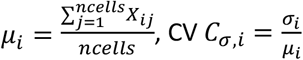, and dropout rate 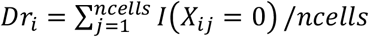, where 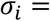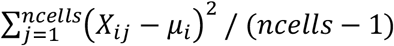 is the standard deviation of a gene, *I* (*X*_*i j*_ = 0) is an indicator function that equals 1 if *X*_*i j*_ = 0 (indicating a dropout event) and 0 otherwise, *i* denotes the gene index and *j* denotes the cell index.

Each list of the three summary statistics is processed independently for each condition under comparison. The list is sorted by the mean expression from small to large, and then a 3D curve containing the cumulative sum of the logarithmic difference between successive elements is built. These differences are defined as follows:

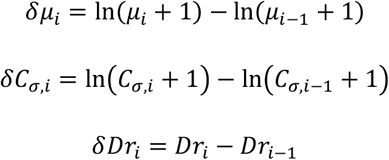

The 3D spline curve contains information of three statistics for every gene, accounting for the variability in the sample. The data-driven curve is built as follows,

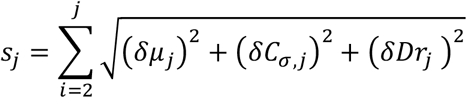

The spline-fit curve provides the “expected” behavior overall statistics of genes in each condition, accounting for intrinsic variability and helping to separate biological variations from the data-driven variation in the curve *s*_*j*_. Formally, gene statistics are in vector form as 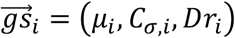, and are compared to the gene values on the 3D spline-fit curve 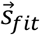, which is described in the same coordinate system. The closest point on the “theoretical” 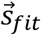, denoted as 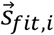 compared to the gene statistics describes how much it deviates from its “expected” behavior: 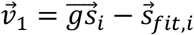 for condition 1, and likewise for condition 2, 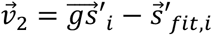, where 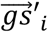 denotes the gene statistics and 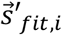 the closest point to the *i*-th gene statistics for the condition 2. 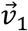 and 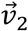 represent the variability of two conditions. The difference in the variability of the two conditions is given as 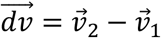 The magnitude of the difference, 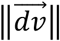, is the DV score. Genes with a significantly greater DV score are considered DV genes. In addition, the difference in magnitude of 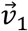 and 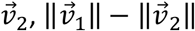 gives a sense of the direction of variability changes between two conditions.

## Code Availability

Code implementing the spline-DV method in both R and MATLAB is available at https://github.com/cailab-tamu/Spline-DV along with example data sets for testing.

## Acknowledgement

We are grateful to Dr. Liang Hu for sharing the hESC-EC scRNA-seq data. We acknowledge the use of advanced computing resources provided by Texas A&M High Performance Research Computing in conducting parts of this research. This study was funded by the U.S. Department of Defense (DoD, GW200026) for J.J.C, Allen Endowed Chair in Nutrition & Chronic Disease Prevention for R.S.C., and the Cancer Prevention & Research Institute of Texas (CPRIT, RP230204) for J.J.C. and R.S.C.

## Author Contributions

**V.G**.: Formal Analysis, Investigation, Visualization, Writing – Original Draft, Writing – Review & Editing. **S.G**.: Methodology, Software, Visualization, Writing – Review & Editing. **S.R**.: Methodology, Software, Visualization, Writing – Review & Editing. **R.C**.: Supervision, Writing – Review & Editing. **J.J.C**.: Conceptualization, Writing – Original Draft, Visualization, Writing – Review & Editing, Project Administration. All authors reviewed and approved the manuscript.

