## Supplementary Information for "Beyond Differential Expression: Embracing Cell-to-Cell Variability in Single-Cell Gene Expression Data Analysis"

**Supplementary Table 1.** Top DV genes from Case Study 2

| Rank | Gene | DV |
|---|---|---|
| 1 | Acta2 | 0.641048 |
| 2 | Gpx3 | 0.543834 |
| 3 | Dpt | 0.520874 |
| 4 | Igfbp4 | 0.448884 |
| 5 | Alcam | 0.398971 |
| 6 | Trf | 0.362375 |
| 7 | Gadd45g | 0.328445 |
| 8 | Spidr | 0.319335 |
| 9 | Col1a1 | 0.319328 |
| 10 | Pim1 | 0.319005 |
| 11 | C3 | 0.3093 |
| 12 | Hspb1 | 0.302271 |
| 13 | Timp1 | 0.298288 |
| 14 | Fam180a | 0.291969 |
| 15 | Creld2 | 0.287111 |
| 16 | Gas6 | 0.283233 |
| 17 | Mdk | 0.281838 |
| 18 | S100a6 | 0.274491 |
| 19 | Inmt | 0.270735 |
| 20 | Serpine2 | 0.267025 |
| 21 | Hspa1a | 0.256424 |
| 22 | Htra3 | 0.234665 |
| 23 | Igf1 | 0.222734 |
| 24 | Myl9 | 0.217734 |
| 25 | Ifit3 | 0.214635 |
| 26 | Clu | 0.211816 |
| 27 | Col1a2 | 0.200576 |
| 28 | Lmcd1 | 0.1999 |
| 29 | Gm12216 | 0.198946 |
| 30 | Atf3 | 0.194491 |
| 31 | Stmn1 | 0.193674 |
| 32 | Tuba1b | 0.192311 |
| 33 | Hk2 | 0.187582 |
| 34 | Mest | 0.183297 |
| 35 | Emp1 | 0.180565 |
| 36 | Hmgb2 | 0.179847 |
| 37 | Nr4a1 | 0.17966 |
| 38 | Hp | 0.178242 |
| 39 | Thbs1 | 0.17351 |
| 40 | Wnt4 | 0.17032 |
| 41 | C4b | 0.169897 |
| 42 | Clec4g | 0.165912 |
| 43 | Ccl7 | 0.165499 |
| 44 | Rgs4 | 0.165233 |
| 45 | Cdkn1a | 0.164468 |
| 46 | Adamts4 | 0.163991 |
| 47 | Wt1 | 0.162702 |
| 48 | Plat | 0.162312 |
| 49 | Aebp1 | 0.15961 |
| 50 | Ifit1 | 0.156609 |
| 51 | Id2 | 0.155293 |
| 52 | Mt2 | 0.154613 |
| 53 | Lpl | 0.15058 |
| 54 | Mmp2 | 0.150488 |
| 55 | Col3a1 | 0.148994 |
| 56 | Hspa1b | 0.148756 |
| 57 | Tagln | 0.148256 |
| 58 | Serpina3g | 0.147691 |
| 59 | Ndufa4l2 | 0.143252 |
| 60 | Gadd45b | 0.142686 |
| 61 | Ccl2 | 0.141078 |
| 62 | Igfbp3 | 0.138713 |
| 63 | Col8a1 | 0.136395 |
| 64 | Cpxm1 | 0.136337 |
| 65 | Egr1 | 0.136025 |
| 66 | Lcn8 | 0.134039 |
| 67 | Loxl1 | 0.133334 |
| 68 | Rsad2 | 0.130822 |
| 69 | Timp3 | 0.128052 |
| 70 | Mgst1 | 0.127589 |
| 71 | Spon2 | 0.124167 |
| 72 | Lox | 0.123773 |
| 73 | Vipr1 | 0.123208 |
| 74 | Fhl2 | 0.121531 |
| 75 | Nbl1 | 0.12069 |
| 76 | Cadm3 | 0.119845 |
| 77 | Col6a3 | 0.118604 |
| 78 | Sdc4 | 0.117904 |
| 79 | Cyp2e1 | 0.117886 |
| 80 | Isg15 | 0.117128 |
| 81 | Tpm2 | 0.113735 |
| 82 | Lum | 0.113198 |
| 83 | Gm13889 | 0.111806 |
| 84 | Fosb | 0.111512 |
| 85 | Angptl6 | 0.111508 |
| 86 | Nupr1 | 0.110457 |
| 87 | Ptgis | 0.109462 |
| 88 | Pdgfra | 0.109425 |
| 89 | Gm2a | 0.109323 |
| 90 | Rgs2 | 0.108889 |
| 91 | Hmgn2 | 0.108813 |
| 92 | Cyp4b1 | 0.108487 |
| 93 | Meg3 | 0.108048 |
| 94 | Shisa2 | 0.104187 |
| 95 | Tnfrsf11b | 0.104024 |
| 96 | Ramp2 | 0.10335 |
| 97 | Wfdc1 | 0.103239 |
| 98 | Slc3a2 | 0.102831 |
| 99 | Eln | 0.102385 |
| 100 | Tubb4b | 0.102293 |
| 101 | Dhrs3 | 0.100836 |
| 102 | Mfap2 | 0.100441 |
| 103 | Gsn | 0.100157 |
| 104 | Actg1 | 0.099799 |
| 105 | Ifit3b | 0.099686 |
| 106 | Cxcl10 | 0.099609 |
| 107 | Tgfb3 | 0.098811 |
| 108 | Capn6 | 0.098574 |
| 109 | Plagl1 | 0.098114 |
| 110 | Itm2a | 0.097898 |
| 111 | Cenpa | 0.097874 |
| 112 | Angptl4 | 0.096967 |
| 113 | Tuba1a | 0.096903 |
| 114 | Ccl19 | 0.095336 |
| 115 | Rgs5 | 0.095318 |
| 116 | Zbp1 | 0.095154 |
| 117 | Cdk1 | 0.094337 |

| Rank | Gene | DV |
| --- | --- | --- |
| 118 | Eno1 | 0.094072 |
| 119 | Gapdh | 0.093669 |
| 120 | Htra1 | 0.093629 |
| 121 | Fbxl22 | 0.093559 |
| 122 | Aard | 0.093002 |
| 123 | Csf1 | 0.092522 |
| 124 | Cebpb | 0.092447 |
| 125 | Trib1 | 0.092319 |
| 126 | Errfi1 | 0.092311 |
| 127 | Ifi27 | 0.091847 |
| 128 | Pth1r | 0.091739 |
| 129 | Calcrl | 0.091136 |
| 130 | Tcf21 | 0.090947 |
| 131 | Dkk3 | 0.09076 |
| 132 | Clec11a | 0.089883 |
| 133 | Nnat | 0.08976 |
| 134 | Egr2 | 0.089058 |
| 135 | Spry1 | 0.088892 |
| 136 | Ugdh | 0.087064 |
| 137 | Maf | 0.086613 |
| 138 | Ier2 | 0.086062 |
| 139 | Fam162a | 0.085875 |
| 140 | Apoc1 | 0.085069 |
| 141 | Fbln1 | 0.084564 |
| 142 | Kcne4 | 0.084434 |
| 143 | Lgals1 | 0.084237 |
| 144 | Itgb3 | 0.08396 |
| 145 | Cxcl14 | 0.08349 |

| Rank | Gene | DV |
| --- | --- | --- |
| 146 | Jun | 0.082866 |
| 147 | Arl6ip1 | 0.082289 |
| 148 | Fabp5 | 0.081992 |
| 149 | Tpm4 | 0.081486 |
| 150 | Bicc1 | 0.080783 |
| 151 | Bhlhe40 | 0.080447 |
| 152 | Nfkbia | 0.080118 |
| 153 | Neat1 | 0.079834 |
| 154 | Socs3 | 0.079682 |
| 155 | Ddit4 | 0.079415 |
| 156 | Ppp1r14a | 0.079178 |
| 157 | Tinagl1 | 0.078112 |
| 158 | Mmp14 | 0.077939 |
| 159 | Cks2 | 0.077789 |
| 160 | Ly6a | 0.077267 |
| 161 | Csrp2 | 0.077231 |
| 162 | Msc | 0.077015 |
| 163 | Ehd3 | 0.075625 |
| 164 | Adamts5 | 0.075129 |
| 165 | Serpine1 | 0.074601 |
| 166 | Mmp23 | 0.074459 |
| 167 | Gbp2 | 0.073995 |
| 168 | Ctsk | 0.073856 |
| 169 | Apold1 | 0.073849 |
| 170 | Cnn2 | 0.073849 |
| 171 | Osmr | 0.073831 |
| 172 | Ntm | 0.073424 |
| 173 | Hgf | 0.073368 |

| Rank | Gene | DV |
| --- | --- | --- |
| 174 | Vcam1 | 0.072792 |
| 175 | Hsp90aa1 | 0.072028 |
| 176 | Cd200r3 | 0.071332 |
| 177 | Nrxn1 | 0.070696 |
| 178 | Dnajb1 | 0.070095 |
| 179 | Hsp90ab1 | 0.069434 |
| 180 | Mustn1 | 0.069282 |
| 181 | Tmem100 | 0.069248 |
| 182 | Itga8 | 0.068807 |
| 183 | Adamts1 | 0.068372 |
| 184 | Lrp4 | 0.068344 |
| 185 | Cryab | 0.068244 |
| 186 | Crip1 | 0.067621 |
| 187 | Sparcl1 | 0.067481 |
| 188 | Tnc | 0.067374 |
| 189 | Tpm1 | 0.067284 |
| 190 | Isyna1 | 0.067135 |
| 191 | Slmap | 0.066619 |
| 192 | Fn1 | 0.065684 |
| 193 | C6 | 0.065462 |
| 194 | Tnfsf9 | 0.065402 |
| 195 | Gm12840 | 0.065368 |
| 196 | Hspa8 | 0.065077 |
| 197 | Plekhf1 | 0.06468 |
| 198 | Myl12a | 0.063833 |
| 199 | Ncam1 | 0.063698 |
| 200 | Derl3 | 0.063447 |

**Supplementary Table 2.** Top DV genes from Case Study 3

| Rank | Gene | DV |
|---|---|---|
| 1 | ANPEP | 0.303824 |
| 2 | DDX60 | 0.263652 |
| 3 | NRXN3 | 0.253701 |
| 4 | LGR5 | 0.243332 |
| 5 | FCGBP | 0.222222 |
| 6 | NEDD4L | 0.215333 |
| 7 | LINC01876 | 0.214317 |
| 8 | XACT | 0.213588 |
| 9 | KCNMA1 | 0.201338 |
| 10 | AGBL4 | 0.190562 |
| 11 | MUC2 | 0.186617 |
| 12 | SELENBP1 | 0.185844 |
| 13 | GUCA2A | 0.178930 |
| 14 | CD55 | 0.177503 |
| 15 | AFF3 | 0.176702 |
| 16 | SMOC2 | 0.171783 |
| 17 | ROBO2 | 0.167587 |
| 18 | ITPR2 | 0.166068 |
| 19 | GPHN | 0.163363 |
| 20 | GDA | 0.159377 |
| 21 | CCSER1 | 0.152209 |
| 22 | NCKAP5 | 0.149180 |
| 23 | NEBL | 0.145820 |
| 24 | PTPRO | 0.144630 |
| 25 | CELF2 | 0.143912 |
| 26 | APBB2 | 0.143358 |
| 27 | NOX1 | 0.141960 |
| 28 | CDH13 | 0.140267 |
| 29 | BCAS1 | 0.139714 |
| 30 | SLC15A1 | 0.137512 |
| 31 | STOX2 | 0.134814 |
| 32 | AGAP1 | 0.134369 |
| 33 | MEF2C | 0.133274 |
| 34 | SAT1 | 0.133217 |
| 35 | CHST11 | 0.127661 |
| 36 | ADAMTSL1 | 0.122507 |
| 37 | ARL15 | 0.121215 |
| 38 | SLC5A1 | 0.120637 |
| 39 | SOX4 | 0.119905 |
| 40 | PCK1 | 0.118032 |
| 41 | FGGY | 0.117755 |
| 42 | HDAC2 | 0.117703 |
| 43 | APOLD1 | 0.117489 |
| 44 | PLCB4 | 0.116975 |
| 45 | ZNF69 | 0.115434 |
| 46 | ODF2L | 0.115276 |
| 47 | CHRNA7 | 0.115091 |
| 48 | KIZ | 0.114827 |
| 49 | LPIN2 | 0.113976 |
| 50 | FER1L6 | 0.113797 |
| 51 | EZR | 0.113768 |
| 52 | EYS | 0.112857 |
| 53 | DIAPH3 | 0.112421 |
| 54 | EEF2 | 0.112304 |
| 55 | PTPRN2 | 0.111063 |
| 56 | CEACAM1 | 0.110761 |
| 57 | NHS | 0.110497 |
| 58 | PDE9A | 0.110220 |
| 59 | PDE3B | 0.107334 |
| 60 | RUBCNL | 0.106414 |
| 61 | HSP90AA1 | 0.106024 |
| 62 | HSPH1 | 0.105079 |
| 63 | BACH2 | 0.103974 |
| 64 | MALRD1 | 0.103869 |
| 65 | CENPP | 0.103806 |
| 66 | MECOM | 0.102554 |
| 67 | FYB1 | 0.102453 |
| 68 | OCLN | 0.102426 |
| 69 | DANT2 | 0.102293 |
| 70 | GMDS | 0.101803 |
| 71 | EBF1 | 0.101428 |
| 72 | NBPF19 | 0.098867 |
| 73 | BIRC3 | 0.098149 |
| 74 | TMEM117 | 0.097744 |
| 75 | SLCO2A1 | 0.096119 |
| 76 | LTBP1 | 0.096039 |
| 77 | PTPRR | 0.095908 |
| 78 | EPS8 | 0.095749 |
| 79 | PPM1L | 0.095594 |
| 80 | SYTL2 | 0.095529 |
| 81 | MS4A1 | 0.094119 |
| 82 | DAB1 | 0.094020 |
| 83 | SSH2 | 0.094002 |
| 84 | MALT1 | 0.093431 |
| 85 | MIR4435-2HG | 0.093347 |
| 86 | TRANK1 | 0.093085 |
| 87 | RNF213 | 0.093082 |
| 88 | MUC12 | 0.092619 |
| 89 | SLC9A2 | 0.092074 |
| 90 | CFTR | 0.091797 |
| 91 | TTN | 0.091356 |
| 92 | HHLA2 | 0.091302 |
| 93 | ANKRD44 | 0.089784 |
| 94 | XDH | 0.089670 |
| 95 | RORA | 0.089641 |
| 96 | NXPE1 | 0.089564 |
| 97 | KAZN | 0.089265 |
| 98 | RUNX1 | 0.089124 |
| 99 | MID1 | 0.088861 |
| 100 | NORAD | 0.088455 |
| 101 | TCF4 | 0.088319 |
| 102 | SLC12A2 | 0.087906 |
| 103 | RNF152 | 0.087201 |
| 104 | CPA6 | 0.086509 |
| 105 | SCMH1 | 0.086487 |
| 106 | ITGA6 | 0.085528 |
| 107 | ANKRD36 | 0.085483 |
| 108 | TACC1 | 0.085303 |
| 109 | PTMA | 0.084799 |
| 110 | CENPF | 0.084608 |
| 111 | IQCM | 0.084572 |
| 112 | PPARG | 0.084511 |
| 113 | LRIG1 | 0.083986 |
| 114 | INPP4B | 0.083971 |
| 115 | ARHGAP15 | 0.082943 |
| 116 | EDA | 0.082640 |

| Rank | Gene | DV |
|---|---|---|
| 117 | CEP112 | 0.082558 |
| 118 | HUNK | 0.082326 |
| 119 | DUSP6 | 0.082125 |
| 120 | FRMD5 | 0.081689 |
| 121 | FERMT1 | 0.080726 |
| 122 | EDAR | 0.080337 |
| 123 | CAMK2N1 | 0.080128 |
| 124 | SGK1 | 0.080096 |
| 125 | FGFR2 | 0.079386 |
| 126 | P3H2 | 0.079177 |
| 127 | RASEF | 0.079125 |
| 128 | PID1 | 0.078912 |
| 129 | MCF2L | 0.078902 |
| 130 | LIPH | 0.078771 |
| 131 | CLIC5 | 0.078727 |
| 132 | JAG1 | 0.078461 |
| 133 | PRKCB | 0.078065 |
| 134 | TMEM132D | 0.077857 |
| 135 | CDH17 | 0.077849 |
| 136 | PFKFB2 | 0.077791 |
| 137 | SLC39A11 | 0.077443 |
| 138 | MACF1 | 0.077249 |
| 139 | SI | 0.076936 |
| 140 | PYGB | 0.076643 |
| 141 | SNTB1 | 0.076357 |
| 142 | GPCPD1 | 0.076252 |
| 143 | PTPRB | 0.076071 |
| 144 | ZMYM2 | 0.076053 |

| Rank | Gene | DV |
|---|---|---|
| 145 | RAPGEF2 | 0.075847 |
| 146 | POLA1 | 0.075617 |
| 147 | ABTB2 | 0.075145 |
| 148 | PPP1R9A | 0.074775 |
| 149 | ATP9A | 0.074654 |
| 150 | USP53 | 0.074642 |
| 151 | CNKSR3 | 0.074567 |
| 152 | KCNK1 | 0.074529 |
| 153 | DPP10 | 0.074195 |
| 154 | SPATS2L | 0.073868 |
| 155 | TBC1D22A | 0.073291 |
| 156 | VAV3 | 0.073136 |
| 157 | SPINK5 | 0.073021 |
| 158 | VIPR1 | 0.072994 |
| 159 | SAMD9 | 0.072927 |
| 160 | TMSB4X | 0.072540 |
| 161 | RFC3 | 0.072510 |
| 162 | JUND | 0.072038 |
| 163 | VTI1A | 0.072024 |
| 164 | DNAJC15 | 0.071958 |
| 165 | NR1H4 | 0.071827 |
| 166 | SPTBN1 | 0.071798 |
| 167 | TRPM6 | 0.071746 |
| 168 | HMGCS2 | 0.071675 |
| 169 | TNRC6B | 0.071615 |
| 170 | ST6GAL1 | 0.071548 |
| 171 | SORBS1 | 0.071509 |
| 172 | MYRIP | 0.071467 |

| Rank | Gene | DV |
|---|---|---|
| 173 | FAT1 | 0.071350 |
| 174 | DPH6 | 0.070893 |
| 175 | PCSK5 | 0.070764 |
| 176 | NFIA | 0.070478 |
| 177 | CRACR2A | 0.070323 |
| 178 | RBPJ | 0.070218 |
| 179 | CA12 | 0.069763 |
| 180 | BRIP1 | 0.069686 |
| 181 | CEP70 | 0.069544 |
| 182 | PRELID2 | 0.068942 |
| 183 | DNAH14 | 0.068705 |
| 184 | OXR1 | 0.068519 |
| 185 | REP15 | 0.068372 |
| 186 | PTPRD | 0.068072 |
| 187 | PRSS23 | 0.067656 |
| 188 | FAM83E | 0.067510 |
| 189 | RAI14 | 0.067352 |
| 190 | NFAT5 | 0.067329 |
| 191 | PRIM2 | 0.067091 |
| 192 | SRI | 0.067053 |
| 193 | GMDS-DT | 0.067024 |
| 194 | LINC02086 | 0.066866 |
| 195 | KDM2A | 0.066557 |
| 196 | SORBS2 | 0.066029 |
| 197 | FSIP2 | 0.065818 |
| 198 | EPB41L3 | 0.065602 |
| 199 | CENPT | 0.065511 |
| 200 | PPARGC1A | 0.065321 |
